# Learning to learn: Single session acquisition of new rules by freely-moving mice

**DOI:** 10.1101/2023.03.28.534599

**Authors:** Amir Levi, Noam Aviv, Eran Stark

**Affiliations:** Department of Physiology and Pharmacology, Faculty of Medicine, Tel Aviv University, Tel Aviv 6997801, Israel; Sagol School of Neuroscience, Tel Aviv University, Tel Aviv 6997801, Israel; Sagol Department of Neurobiology, Haifa University, Haifa 3103301, Israel

## Abstract

Learning from examples and adapting to new rules are fundamental attributes of human cognition. However, it is unclear what conditions allow for fast and successful learning, especially in non-human subjects. To determine how rapidly freely-moving mice can learn a new rule, we designed a two-alternative forced-choice visual discrimination paradigm in which the rules governing the task can change between sessions. We find that experienced animals can learn a new rule after being exposed to only five training and three testing trials. The propensity for single session learning improves over time and is accurately predicted based on animal experience and rule difficulty. After establishing procedural learning of a paradigm, mice continuously improve their performance on new rules. Thus, mice learn to learn.

## Introduction

Solving a discrimination task for a reward involves associative operant learning^1^. Through experience, we learn the relationship between stimuli, actions, and results, aligning our actions with the desired outcomes. As we practice the task, certain aspects become easier and are eventually performed implicitly. Now consider a sudden change in some of the rules of a well-known task. The ability to adapt to new rules or learn from a small training set is a fundamental attribute of human cognition^2^. However, it is not clear which behavioral patterns underlie adaptation, and what conditions allow fast and successful learning of new rules governing a well-known task.

One way to learn a new rule governing a well-known discrimination task is to generalize from a known rule. Rodents, which provide a convenient model system for studying the neurobiological basis of behavior, can use generalization^3^ and transfer^4,5^ to learn new rules. However, generalization cannot be used when the new rule is uncorrelated with the previously learned rules. A second option for successful performance when the rules change is categorization, an established ability among rodents^6,7^. However, new rules do not necessarily fall into previously acquainted categories. Third, higher levels of attention and experience may facilitate faster learning^8^. Attending to details increases learning rate^9^. Furthermore, because learning is never carried out on a completely blank slate, previous knowledge (i.e., experience) may facilitate the acquisition of new rules via learning sets^10^ and schemas^11,12^. Indeed, during repeated changes to the rules of a well-known task, experience may facilitate learning.

Although advantageous, extensive experience is not necessary for learning. Fast and even one-shot learning^13^ were previously demonstrated in laboratory rodents. During fear conditioning, a stimulus is associated with a single exposure to an aversive experience, leading to avoidance learning^14,15^. Rodents also excel in spatial learning^16^ and can learn from a few exposures^17,18^ or even from a single spatial experience^19–21^. However, classical learning and operant learning are associated with distinct neuronal mechanisms^22,23^. Furthermore, although rodents are capable of quick learning of naturalistic tasks, many laboratory operant learning tasks are conducted on naïve subjects^12^ and take weeks to learn^6,24–28^. Thus, it is unknown how rapidly can rodents learn a new set of associations within a well-known setting and what is the specific contribution of experience.

Using a fully automated two-alternative forced-choice (2AFC) paradigm, we find that every mouse learns a new visual discrimination rule in a single session. When single session learning (SSL) occurs, mice perform the task successfully after being exposed to only three testing trials. Physically difficult rules are less likely to result in SSL, and experienced mice are more likely to achieve SSL. SSL can be achieved for particularly difficult rules if conditions for generalization from a previously learned rule are favorable. Thus, mice learn to learn.

## Results

### All mice learn at least one new visual rule within a single session

We designed a fully automated 2AFC discrimination paradigm for freely-moving mice in a T-maze (**Fig. 1A**). All subjects were hybrid mice (first-generation offspring of FVB/NJ females and C57BL/6J males^21^; n=6) that were pretrained (“shaped”) over several days (median [interquartile range, IQR]: 5 [4 5] days). To maximize learning rate and minimize frustration, every post-shaping session was divided into blocks (**Fig. 1B**). Each block consisted of five training trials followed by ten testing trials. During training trials, the subjects were presented with the same stimuli as during testing trials, but the correct choice was enforced by opening only one door at the T-junction (**Fig. 1C**). Mice received a water reward after every training trial and every successful testing trial.

**Figure 1.**
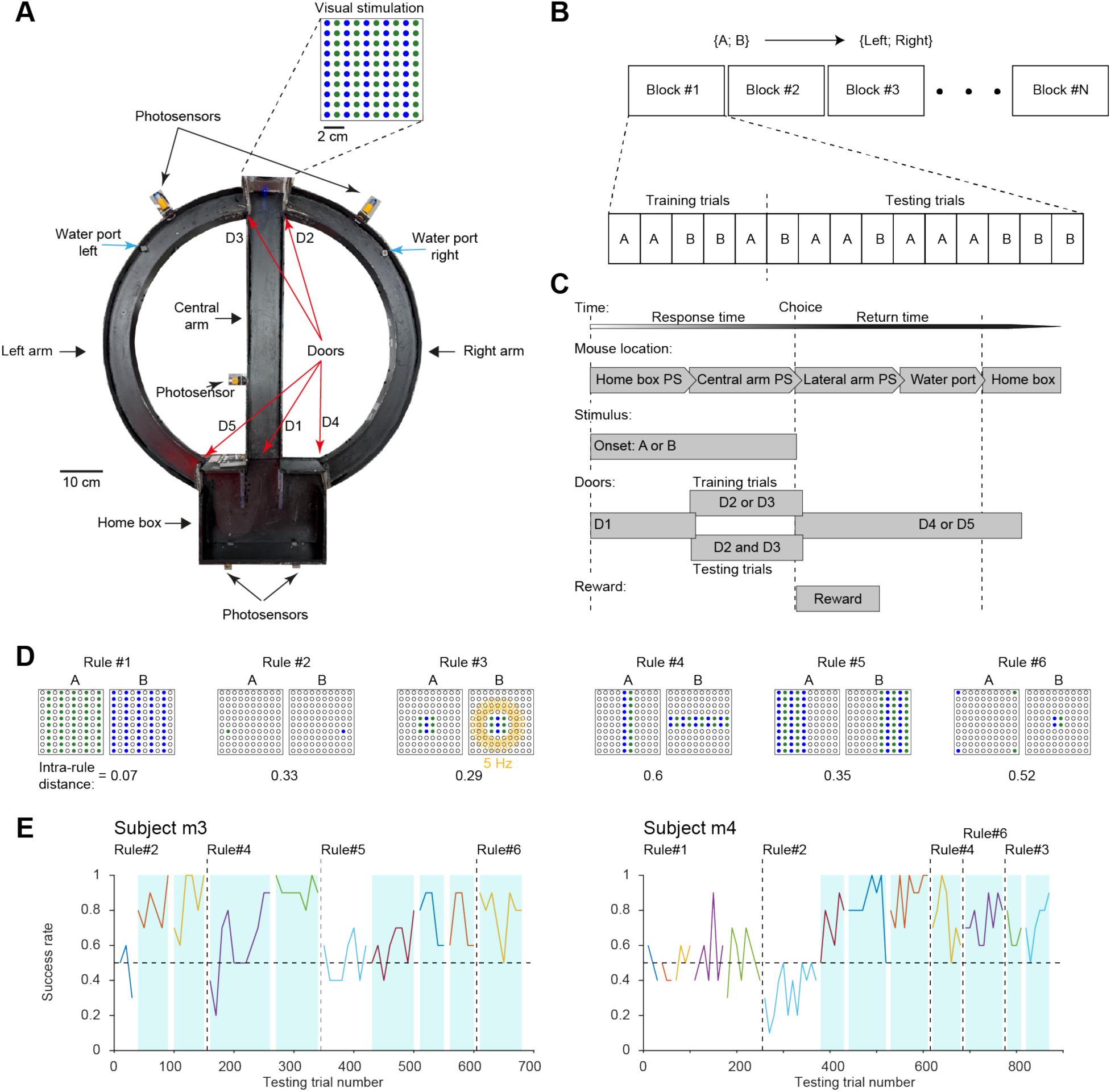
All mice learn at least one new visual rule within a single session. (**A**) The fully automated apparatus used to train freely-moving mice on a 2AFC discrimination task. Successful trials are reinforced by water. **Top**, The display used to generate visual rules, located at the far end of the central arm, consisting of ten alternating columns of green and blue LEDs. (**B**) Session structure. Every session is composed of blocks, each consisting of five training trials followed by ten testing trials. (**C**) Trial structure. During training trials, only the correct choice is available to the animal since only one door at the T-junction is open (D2 or D3). (**D**) Six visual rules are used to govern the task. On every session only one rule is used. **Bottom**, The physical properties of two visual stimuli {A; B} are used to derive an intra-rule distance metric, quantifying rule difficulty. (**E**) Success rate as a function of testing trial number for two example subjects, m3 and m4. Sessions are represented by distinct colors, and success rate is averaged per block. A blue background highlights successful sessions (p<0.05, Binomial test comparing to chance level, 0.5).

During every session, a display of 100 green and blue LEDs (**Fig. 1A, inset**) was used to provide two distinct visual stimuli {A; B} for governing the task. Thus, a rule was composed of two stimulus-response associations, an affinity between stimulus A and “go left”, and an affinity between B and “go right”. Only one rule was employed during a given session. The physical properties of the two visual stimuli allowed characterizing every rule by an intra-rule distance that ranges [0,1] and is smaller for more difficult rules (**Fig. 1D**). For example, rule#1 consisted of {stimulus A: all green LEDs are on; stimulus B: all blue LEDs are on} and had an intra-rule distance of 0.07, indicating the rule is relatively difficult.

A session was denoted as successful if performance was consistently above chance (0.5; p<0.05, Binomial test). A rule was deemed successfully learned if a successful session was performed while the task was governed by the rule. Using the six-rule pool (**Fig. 1D**), the six mice were tested on 28 rules, with every subject exposed to a median [range] of 5 [3,6] rules. All subjects achieved SSL of at least one rule. When exposed to a new rule, a median [IQR] of 90 [70 138] testing trials were carried out until a successful session was completed (**Fig. S1B-C**). For example, after pretraining (shaping) on rule#2, subject m3 performed the task successfully during the second and third testing sessions (**Fig. 1E, left**). The rule was then changed to rule#4, which was learned in a single session. m3 was tested on three new rules and achieved SSL on two. Similar results were obtained for all subjects, with a median [range] of 2 [1,3] SSL rules per subject (**Table 1**; **Fig. S1**). Thus, all mice learned a least one new visual rule within a single session.

**Table 1.**
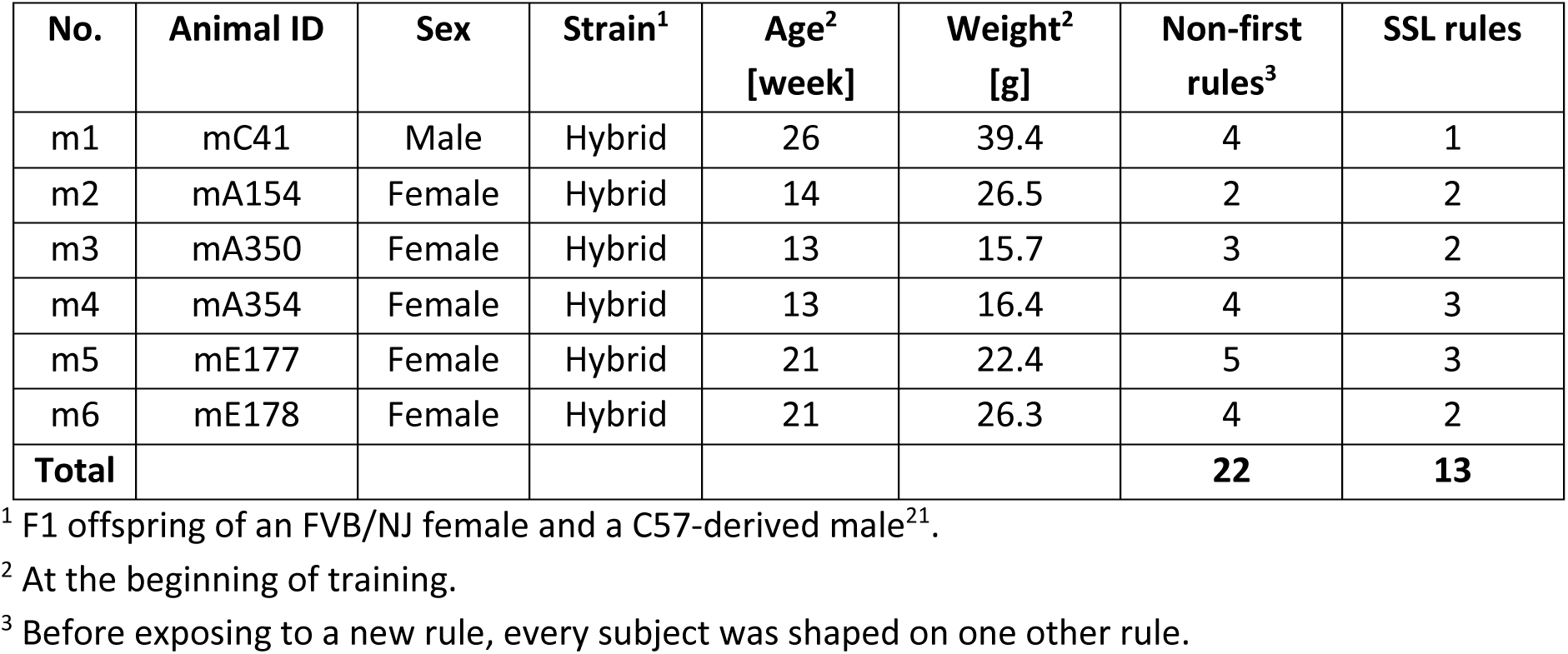
Single session learning in every experimental animal.

### During SSL, mice achieve high success rates from the first trials of the first block

To determine the dynamics of single session learning, we classified sessions according to the learning process and characterized the success rate as a function of block number. Each session was associated with either SSL (e.g., m4, rule#6; **Fig. 1E, right**; **Fig. 2A**), multi-session learning (MSL; e.g., m4, rule#2; **Fig. 1E, right**; **Fig. 2B**), or was unlabeled (e.g., m4, rule#1). By definition, a rule associated with MSL requires more than one session to learn. Since every mouse was pretrained on the first rule encountered by the subject, all first-rule sessions were unlabeled. Of the newly-encountered rules, 13/22 (59%) were SSL, 8/22 (36%; spanning 19 sessions) were MSL, and in one case the rule was not learned (1/22, 5%; m5, rule#1; **Fig. S1**). Thus, in 95% of the cases, mice learn a completely new rule within one to three sessions. SSL success rates (0.7 [0.6 0.8]; n=110 blocks) were higher than MSL success rates (0.6 [0.5 0.7]; n=154; p<0.001, U-test; **Fig. S1E**). To quantify differences in the learning process, a linear model was fit to every SSL and MSL learning curve (**Fig. 2AB**, dashed lines). During MSL, mice exhibited gradually increasing success rates (median slope: 0.013 improvement per block; n=8 rules; p=0.008, Wilcoxon’s test comparing to a zero-slope null; **Fig. 2C, top left**), indicating a learning process. MSL initial success rates were not different from chance (0.5; median: 0.46; p=0.055; **Fig. 2C, bottom left**). In contrast, SSL initial success rates were already above chance (0.75; n=13; p=0.003), and did not increase consistently over blocks (median slope: 0.007; p=0.5; **Fig. 2C, right**). Thus, when SSL occurs, the trials experienced during the first block appear to suffice for learning the new rule.

**Figure 2.**
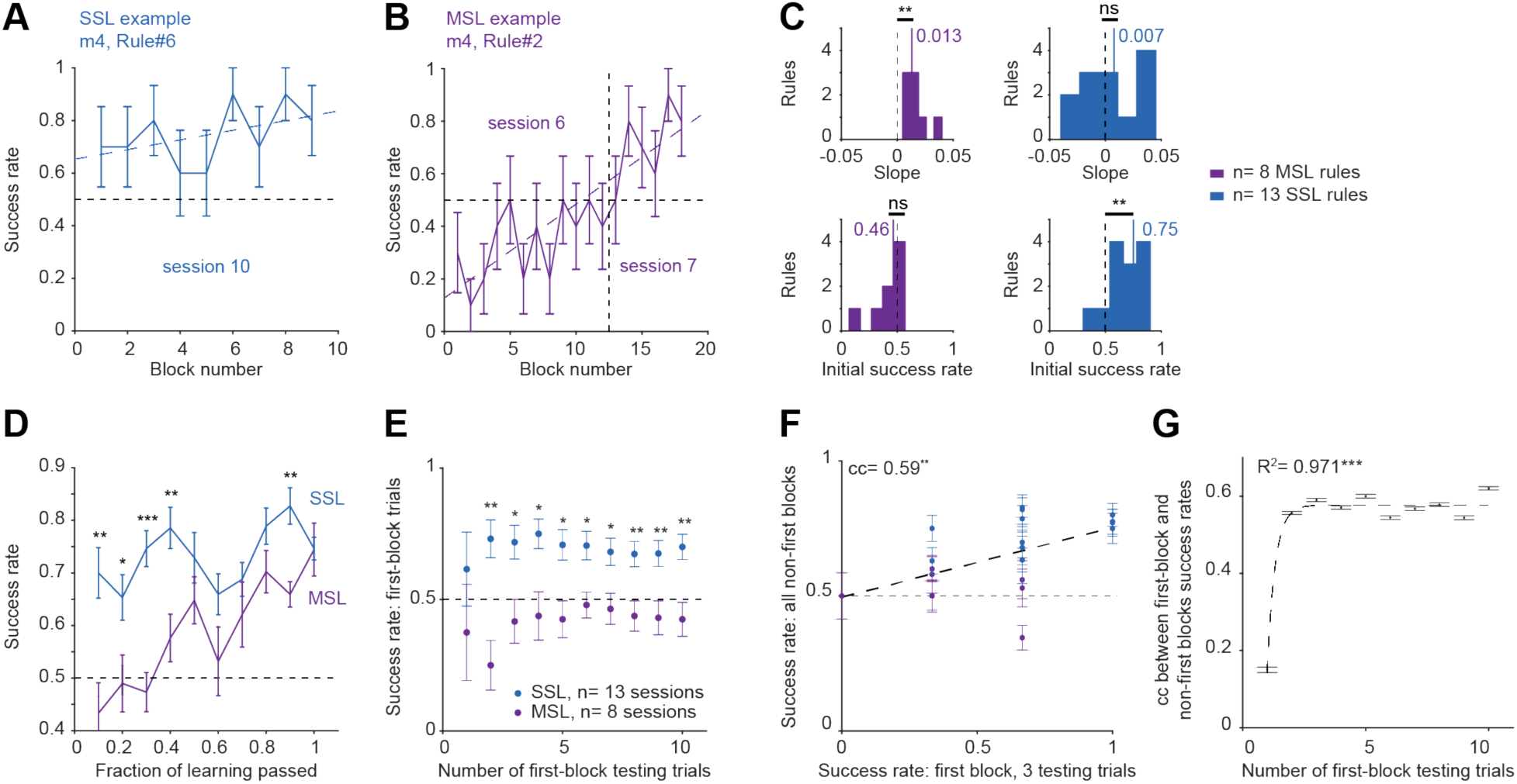
During SSL, mice achieve high success rates from the first trials of the first block. (**A**) Example SSL curve. Here and in **B**, dashed lines represent linear model fit. Here and in BDEF, error bars, SEM. (**B**) Example MSL curve, linear model fitted to the combined two sessions. (**C**) A linear model was fit to every learning curve. **Top**, Slopes. MSL but not SSL curves have slopes consistently above zero. **Bottom**, Intercepts. SSL but not MSL curves have initial success rates consistently above chance. ns/**: p>0.05/p<0.01, Wilcoxon’s test comparing to chance level. (**D**) Success rate of SSL is consistently higher compared with MSL during the initial 40% of the session. Here and in **E**, */**/***: p<0.05/p<0.01/p<0.001, U-test. (**E**) Success rates of the first one, two, three… or ten testing trials of SSL and MSL sessions. (**F**) Success rates in all non-first blocks of the first sessions of newly encountered rules vs. success rates in the first 3 trials of the session. Dashed line, linear fit. cc, rank correlation coefficient; **: p<0.01, permutation test. (**G**) cc-s between success rate during the first few first-block testing trials and all non-first blocks, plotted against the number of first-block testing trials. Dashed line, exponential fit. ***, p<0.001, F-test. Error bars, SD. The correlation stabilizes after three testing trials.

Linear models do not necessarily capture differences between learning curves. Instead of considering models that require more free parameters (e.g., sigmoid), we time-warped every curve to unity duration and averaged all learning curves of a given type (SSL or MSL; **Fig. 2D**). SSL and MSL success rates during the initial four tenths were higher for SSL compared with MSL sessions (geometric mean, p=0.004, U-test; **Fig. 2D**). The observations complement the linear model fit results showing that during SSL, animals learn the new rule during the very first block.

To understand how subjects can perform above chance from the first block when exposed to a completely new rule, we considered two alternatives. One possibility is that the mice already learn the new rule during the five training trials (**Fig. 1C**) at the beginning of the block. Alternatively, one or more testing trials are required. We found that success rates during the first two (or more) testing trials of a new SSL rule were consistently higher than success rates of first trials in an MSL rule (p<0.05 in all cases; n=13 SSL and 8 MSL first-rule sessions; **Fig. 2E**). To estimate the number of testing trials actually required for learning a new rule, we calculated the correlation between the success rate during the first few testing trials of the first new-rule block, and the success rate during all other same-session blocks (**Fig. 2FG**). There was no consistent correlation when a single first-block trial was considered (cc: 0.15; n=21 sessions; p=0.51, permutation test). However, when two or more first-block trials were considered, the correlation was high (range, [0.54, 0.62]; p<0.05 in all cases; **Fig. 2FG**). An exponential model provided a fit to the correlation as a function of the number of first-block trials (R^2^=0.97; n=10; p<0.001, F-test; **Fig. 2G**), indicating that the correlation already converged for three testing trials (**Fig. 2F**). Thus, after to the five training trials, the first three testing trials exhibit the same correlation as the entire first block with the session success rate, indicating that when the conditions are appropriate, three testing trials suffice for learning a new rule.

### Single session learning is predicted from experience and rule difficulty

What are the physical and mental conditions appropriate for SSL? To assess what determines SSL, we first examined how rule identity affects success rates. Mice exhibited different success rates on different rules (**Fig. 3A**). For instance, rule#6 (**Fig. 1D**) was associated with the highest success (0.8 [0.7 0.9]; n=65 blocks; geometric mean of five comparisons, p=0.00007, Kruskal-Wallis test; **Fig. 3A**). In contrast, rule#1 was associated with the lowest success rates (0.6 [0.5 0.7]; n=67; p=0.006; **Fig. 3A**). Thus, success rates depend on the specific rule.

**Figure 3.**
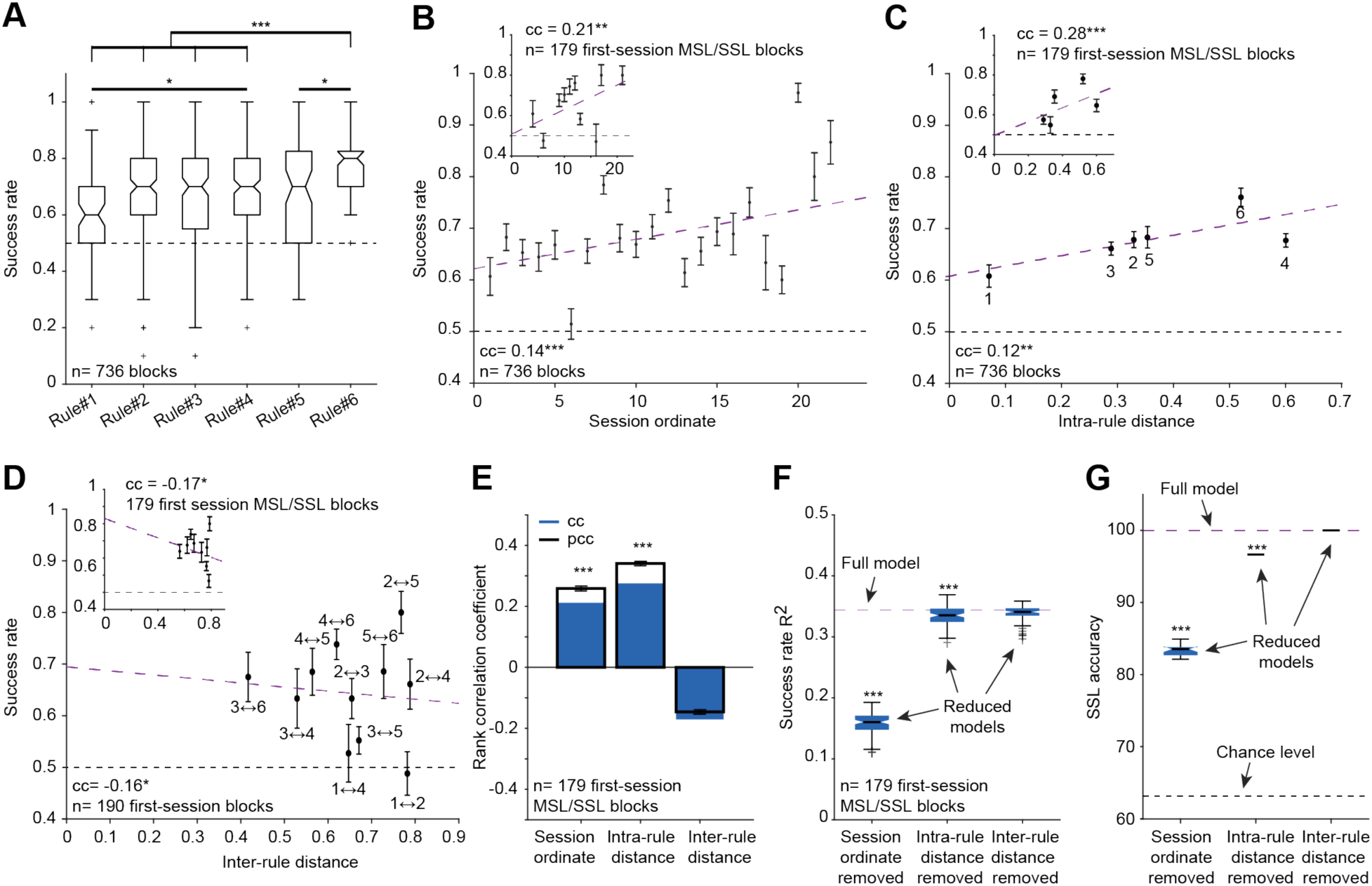
Single session learning is predicted from experience and rule difficulty. (**A**) Success rates for individual rules. */***: p<0.05/p<0.001, Kruskal-Wallis test, corrected for multiple comparisons. Every box plot shows median and IQR, whiskers extend for 1.5 times the IQR in every direction, and a plus indicates an outlier. (**B**) Success rates vs. animal experience, quantified by the session ordinate. Here and in **C** and **D**: purple lines, linear model fit; error bars, SEM. **Inset**, Success rate vs. session ordinate for a reduced set of SSL and MSL sessions. Here and in **C**, **D** and **E**, */**/***: p<0.05/p<0.01/p<0.001, permutation test. (**C**) Success rates vs. rule difficulty, quantified by the intra-rule distance metric (Fig. 1D). (**D**) Success rates vs. the similarity between the new and the previous rule. Only first post-transition sessions are included. (**E**) cc-s and partial rank correlation coefficients (pcc-s) between success rate and the features described in **B**-**D**. Error bars, SD. (**F**) Variance in block success rate (R^2^) explained by cross-validated support vector regression models. ***: p<0.001, Bonferroni-corrected U-test between the R^2^ of the full model and the R^2^ of the reduced model obtained after removing one feature. (**G**) Accuracy in predicting SSL using cross-validated support vector classification. ***: p<0.001, Bonferroni-corrected U-test between the accuracy of the full and reduced models.

When a new session starts and the rule is replaced, mice are compelled to learn the new associations quickly to maximize the reward, requiring memorization of the outcomes of previous trials during the same session (reference memory^29–32^). However, other aspects of the tasks may be learned gradually over multiple sessions. We found that success rates correlated with the accumulated experience of the animal, quantified by the session ordinate (number of previous sessions; cc: 0.14; n=736 blocks; p<0.001, permutation test; **Fig. 3B**). Correlation was not consistently different (p=0.22, bootstrap test) when only the first sessions involving SSL and MSL were included (cc: 0.21; n=179 first-session blocks; p=0.0043; **Fig. 3B, inset**). Thus, when mice are more experienced, performance is better when encountering a new rule.

Even if experience contributes to successful performance, rule difficulty may influence success rate. However, it is not a priori clear what a mouse considers difficult. We quantified rule difficulty using the intra-rule distance metric based on the achromatic physical properties of the stimuli (**Materials and Methods**; **Fig. 1D**). Success rates correlated with intra-rule distance (cc: 0.12; n=736 blocks; p=0.002, permutation test; **Fig. 3C**), and higher correlation (p=0.014, bootstrap test) was observed when only the first SSL and MSL sessions were considered (cc: 0.28; n=179 blocks; p<0.001; **Fig. 3C, inset**). Thus, success rates are correlated with intra-rule distance, especially when a new rule is encountered.

Another possible route to successful performance is generalization from a previously-learned rule. If mice generalize, higher similarity between consecutively-presented rules may yield better performance, and grossly distinct rules may induce confusion. We quantified the similarity between every two rules using the inter-rule distance metric, which is near zero when rules are very similar and near one when rules are very different from one another. In first-session blocks, success rate was anti-correlated with inter-rule distance (cc: −0.16; n=190 first-session blocks; p=0.03, permutation test; **Fig. 3D**). Similar results (p=0.48, bootstrap test) were obtained when only MSL and SSL first-session blocks were considered (i.e., without the 11 blocks of the never-learned rule#1 in m5; cc: −0.17; p=0.023; n=179, **Fig. 3D, inset**). Thus, when a new rule is more similar to the previously-learned rule, success during the first session is already higher.

Based on the foregoing, success level depends on several inter-correlated features (**Fig. S2B**), including session ordinate (**Fig. 3B**), rule difficulty (**Fig. 3C**), inter-rule similarity (**Fig. 3D**), and the success during the initial testing trials (**Fig. 2E-G**). First, we used correlation analysis to disambiguate the features. When considering the first three features, the partial rank correlation coefficient (pcc) was consistently distinct from zero for the session ordinate (pcc: 0.26; n=179 first-session blocks during MSL and SSL; p<0.001, permutation test; **Fig. 3E**) and for intra-rule distance (pcc: 0.34; p<0.001), but not for inter-rule distance (pcc: −0.15; p=0.052). Similar results were obtained when success in the first three first-block trials was used as a fourth feature (**Fig. S2C**).

Second, to determine the total variability of success rate explained by the three features, we used cross-validated support vector regression. Over a third of the variability in block success rate was explained (R^2^=0.34 [0.33 0.35]; 179 blocks; median [IQR] of n=20 independent ten-fold cross-validated models; **Fig. 3F**). By excluding one feature at a time, we found that session ordinate made the dominant contribution to success rate (R^2^=0.16; p<0.001, U-test corrected for multiple comparisons; **Fig. 3F**). Intra-rule distance made a consistent contribution (R^2^=0.33; p<0.001), whereas inter-rule distance did not (R^2^=0.34, p=0.06). Session ordinate made the dominant contribution also when success in the first three first-block trials was included (**Fig. S2D**). Thus, the most important single feature for determining success rate during the first session of a new rule is the accumulated experience.

Knowing that success rate during the first session of a newly encountered rule depends on the accumulated experience and rule difficulty, we assessed what determines SSL. We used cross-validated binary classification (support vector machines) to predict whether a given first session block is SSL or MSL (n=179 blocks). The prediction of SSL from animal experience and physical rule properties was perfect (100% accuracy; **Fig. 3G**). By again excluding one feature at a time, we found that prediction depended on session ordinate (experience; 83% [83% 84%]; p<0.001, U-test; **Fig. 3G**) and on intra-rule distance (rule difficulty; 97% [97% 97%]; p<0.001). Prediction did not depend on inter-rule distance (p=1). A priori, it is possible that if the animal randomly guesses the correct response in the first testing trials, SSL is more likely to be achieved. However, we found that when success in the first three first-block trials was considered, similar results were obtained, with the session ordinate making the dominant contribution in all cases (**Fig. S2EF**). Thus, knowing how experienced an animal is and what are the physical properties of a new rule allows predicting whether SSL will occur.

### When conditions are favorable, mice can generalize from similar yet easier rules

The negligible reduction in variability explained by inter-rule similarity during SSL and MSL sessions (**Fig. 3F**; **Fig. S2D**) suggests that mouse strategy was not based on generalization or categorization according to the previously-learned rule. However, minimizing the usage of generalization may be specific to the set of arbitrarily rules (**Fig. 1D**), for which median [IQR] inter-rule distances were 0.65 [0.53 0.77] (n=11 transitions; **Fig. 3D**). To determine whether mice employ a different strategy when consecutive rules are similar, we tested two of the subjects on a set of five new non-arbitrary rules (m5 and m6; **Fig. 4**). The first new rule was the easiest (highest intra-rule similarity), followed by gradually more difficult rules (**Fig. 4A**). The inter-rule distance between every pair of consecutively-presented rules was 0.04. To minimize the direct effect of the most recently-learned arbitrary rules, animals were kept away from the task for seven days before being presented with the first new rule, and no pretraining (shaping) was conducted. After reaching stable performance on the first new rule (#7), every other rule was presented for one session only. None of the mice managed to learn the first rule during the first session. However, despite the increasing difficulty, both mice achieved SSL for rules #8-10 (**Fig. 4B**). For instance, intra-rule distances of rule#8 and rule#9 were 0.2 and 0.12 (**Fig. 4A**), more difficult than rules #2-6 that were associated with SSL (range: [0.29,0.6]; **Fig. 1D**). Nevertheless, SSL was readily achieved for rules #8-10. Even the most difficult rule (#11; intra-rule distance, 0.04) was associated with SSL in one of the two subjects (m6; **Fig. 4B**). Thus, when inter-rule similarity is high, SSL can be achieved for difficult rules.

**Figure 4.**
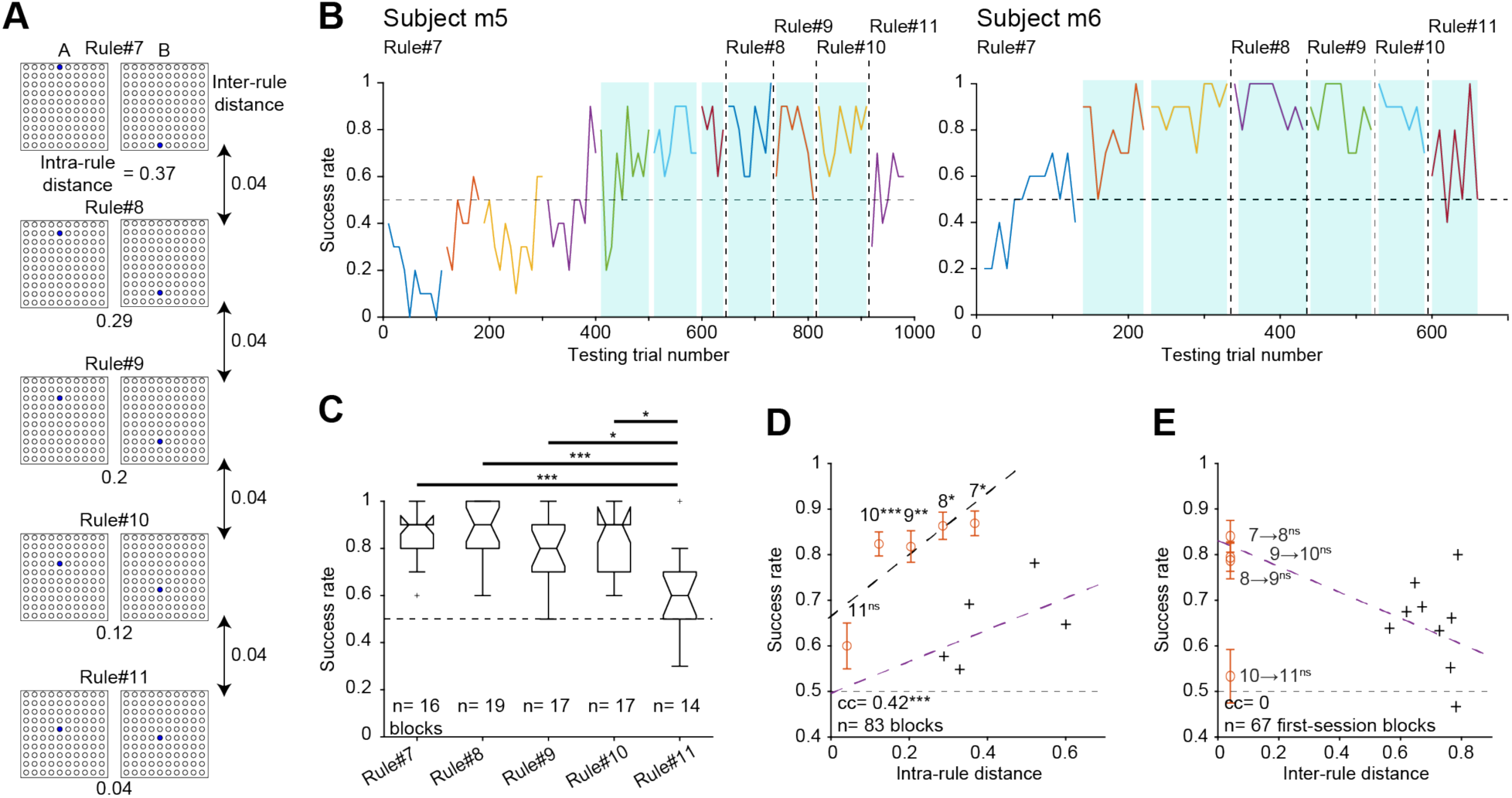
When conditions are favorable, mice can generalize from similar yet easier rules. (**A**) Five visual rules of increasing difficulty distinct from rules #1-6 (Fig. 1D) used consecutively in subjects m5 and m6. (**B**) Success rates as a function of testing trial number. After performance on rule#7 stabilized, a new rule of higher difficulty was employed on every session. All conventions are the same as in Fig. 1E. (**C**) Success rates for individual rules. Here and in **DE**, only the last session of rule#7 was used. */***: p<0.05/p<0.001, Kruskal-Wallis test. (**D**) Success rate vs. rule difficulty. ***: p<0.001, permutation test. Here (and in **E**) crosses and purple lines represent data points and linear models copied from Fig. 3C **inset** (and from Fig. 3D **inset**). Here and in **E**, ns/*/**/***: p>0.05/p<0.05/p<0.01/p<0.001, U-test; error bars, SEM. (**E**) Success rate vs. the similarity between the new and the previous rule.

The set of new rules #7-11 was presented in the same order to both mice. However, success rates during rules #8-10 were not consistently lower than during the last session of rule#7 (n=19, n=17 and n=17 blocks; p=1, p=0.86, and p=0.85, Kruskal-Wallis test; **Fig. 4C**). In contrast, success rates during rule#11 were lower than during rule#10 (p=0.02), although the inter-rule distance between rule#10 and rule#11 was identical to the distance between every other pair of consecutively-presented rules. Success rates were correlated with intra-rule distance (cc: 0.42; n=83 blocks; p<0.001, permutation test; black dashed line in **Fig. 4D**). However, the distances between the points representing rules #7-10 and the linear fit to the arbitrary rules (**Fig. 4D**, purple line) were higher than for rules #1-6 (n=19, n=17, n=17, and n=14 blocks; p=0.013, p=0.0013, p=0.0013, and p=0.0004, U-test; **Fig. 4D**). In contrast, rule#11 did not deviate from the line (n=14 blocks; p=1), suggesting that under the conditions of the task, rule#11 was close to the “just noticeable difference” of visual discrimination. Regardless, success rates of rules #7-10 are higher than expected given the difficulty of the rules and the pattern established by the arbitrary rules (**Fig. 3**), indicating a strategy shift.

The higher-than-expected success rates of rules #8-10 may be explained by generalization from similar yet easier rules or by categorizing the new samples according to an overarching rule, e.g., above/below the horizon. Since all inter-rule distances were identical (0.04), success rates could not be directly correlated with inter-rule distances (**Fig. 4E**). The distances of rules #8-11 from the linear fit to the arbitrary rules were not consistently different than for rules #1-6 (p=0.6, p=0.62, p=0.11, and p=0.056, U-test; **Fig. 4E**). Thus, SSL of particularly difficult rules can be facilitated by generalization from similar yet easier rules. Indeed, success rates during SSL sessions with low inter-rule distances (rules #7-11; 0.85 [0.7 0.9]; n=67 blocks) were consistently higher than for high inter-rule distances (rules #1-6; 0.7 [0.6 0.8], n=112 first-session SSL blocks; p=0.0002, U-test). In sum, when conditions for generalization are more favorable, mice perform SSL with higher success rate.

## Discussion

We tested the ability of mice to learn new rules that span a range of difficulties and quantified the requisite conditions. After acquiring the basic paradigm, subjects gradually improve performance on a new visual rule, and eventually learn a new rule within a single session. Remarkably, all animals achieve SSL of at least one rule. When SSL occurs, experienced mice successfully perform the task after being exposed to five training and three testing trials. SSL is achieved when mice are experienced, when the rule is relatively easy, or when consecutive rules are similar.

### Learning a discrimination rule

We defined a discrimination rule using a pair of stimulus-response associations. In a 2AFC paradigm, the subject can achieve perfect success without learning both associations. When only a single association is learned, solving the task still requires discrimination between the two stimuli. Thus, there are three possibilities for learning the discrimination rules: learn both associations; learn one association; or learn the other association. Yet in all three cases, the discrimination between the two stimuli must be learned.

### Framework for rule learning as part of multilevel learning

Rule learning in a discrimination task with changing rules can be divided into four distinct levels of learning. The first is a type of procedural learning, in which the general task logic is acquired. In the present 2AFC paradigm, the logic involved learning that certain running directions are permitted, that doors open and close, that water may be available at two specific locations, and so on. The logic is learned during the shaping period, in the days prior to the first testing session, is independent of the existence of any specific discrimination rule, and remains relevant for all rules.

The second level involves learning that there is a stimulus-response (S-R) contingency (e.g., stimuli {A; B} are associated with {go left; go right} responses). Here, the second level involved learning that during testing trials, water are available only if specific stimulus-response contingencies are met. The first two levels of learning generate the long-term memory component of standard successive conditional discrimination tasks^24^. When shaping is conducted prior to training on a specific S-R contingency, the two levels may be separated^6,33^. Then, performance is tested on the second level.

The third level involves learning a specific new rule in a well-known setting and is only rarely assessed in animal studies (but see^4,11,12^). Successful performance may be achieved by generalization or transfer from a similar rule^4,5^, by applying a familiar rule to a new set of stimuli (categorization^6,34,35^,), or by associative learning of the new S-R contingencies (the present work). By definition, learning a new rule during a single session requires reference memory^29–32^. If a new rule is not introduced, the third level cannot be assessed. The fourth level, learning that the rules of the task can change between sessions, is a long-term memory component that improves with experience and is independent of the first two levels of learning. In tasks where the rule is fixed during all sessions, the subject may still improve between sessions. However, only when the rules change, the ability of learning to learn a new rule can be assessed. In the present work, we focused on the third and fourth levels.

Tasks that involve generalization^3,34,36^, transfer^4,5^, or categorization^6,34,35^ involve the first two levels but do not require associative learning. The finding that in experienced mice, a new rule can be learned by the end of the first three testing trials indicates that mice can learn to discriminate from a small number of samples, emphasizing the importance of reference memory when conditions in a familiar environment change. Previous work showed that rodents can learn from a few samples in various settings including the Morris water maze^19–21^, fear conditioning^14,15^, and labyrinth navigation^17^. Yet in all aforementioned studies, only two levels of learning (procedural and S-R contingency) were tested. In the radial arm maze^29^ the specific set of arms baited during the session allows to also test the third level (reference memory). However, the rule governing the task remains unchanged. Thus, to the best of our knowledge and with a notable exception^10^, the process of “learning to learn” was not assessed by previous work with animal subjects.

The finding that experience is crucial for SSL becomes clear considering the multilevel learning framework suggested here. The “learning to learn” process (fourth level) implies that a more experienced animal is more flexible to changes in the environment and more likely to learn a new rule quickly. Nevertheless, learning a rule during SSL is by definition limited to a single session. Thus, learning a specific new rule (the third level) is a process distinct from all processes that depend on long-term memory. The fact that easier rules are more likely to be learned suggests a dissociation between the third and fourth levels, supporting the multilevel framework.

### Generalization and categorization

Other cognitive processes may be utilized when two consecutive rules are similar. Learning a new rule when the conditions for generalization are favorable is not independent of the past. Previous work found that rats can use transfer learning to perform a difficult discrimination task^4^. Rodents also excel in categorization^6,37,38^. In both cases, the animal generalizes from previously learned rules and can solve the task without learning new associations, as in the fourth level of learning suggested above. Indeed, when inter-rule distances were lower, success rates were higher. Furthermore, SSL was achieved even for difficult new rules, suggesting that mice did not necessarily learn the new rule, which would have required associative learning and reference memory. Instead, the animals may have categorized the stimuli comprising the new rule using previously acquired knowledge.

### Limitations

While intra-rule distances spanned the entire range of possible values, inter-rule distances were sampled only for high (>0.4) or very low (0.04) values. When encountering intermediate inter-rule distances mice may exhibit yet a third strategy – or change their strategy dynamically according to the relation between intra- and inter-rule distances.

### A neural hypothesis for rule learning

Due to high reproductive rate, genetic control, and well-established tasks for a plethora of behaviors, mice have emerged as a robust model for studying various neuronal mechanisms^39–43^. Head-fixed and freely-moving mice allow combining behavioral and neuronal recordings^24,26^ and manipulations^42–45^. Extensive correlative evidence links neuronal activity with learning, perception, and discrimination^46–48^. Indeed, neuronal mechanisms underlying operant learning are often studied using rodents performing a sensory discrimination task^26,48–52^. The mechanisms that underlie learning in a multi-rule environment have been studied from the theoretical and empirical perspectives^53,54^. However, learning across multiple sessions limits the interpretational power yielded by many rodent learning tasks because in many cases, the same neuronal population cannot be guaranteed to be followed over long durations. Thus, SSL is expected to facilitate the study of the neurophysiological basis of rapid discrimination learning by increasing the overlap between the recorded neuronal activity and the act of learning. Finally, the cellular-network mechanisms underlying the process of learning to learn are unknown. We hypothesize that the framework of multilevel learning will be useful for deciphering these mechanisms.

## Acknowledgements

We thank Ortal Amber-Vitos, Amir Globerson, and Shirly Someck for discussions; Yuval Te’eni for help training animals; and Lirit Levi, Shir Mendelbaum, Pamela Reinagel, Massimo Scanziani, and Hadas E. Sloin for constructive comments. This work was supported by the United States-Israel Binational Science Foundation grant 2015577 (to E.S.); by a European Research Council grant 679253 (to E.S.); by the Israel Science Foundation FIRST Program grant 1871/17 (to E.S.); by the Canadian Institutes of Health Research (CIHR), the International Development Research Centre (IDRC), the Israel Science Foundation (ISF), and the Azrieli Foundation grant 2558/18 (to E.S.); and by the Zimin Institute (to E.S.).

## Author contributions

E.S. conceived the project. A.L. and E.S. designed the apparatus and the experiments. A.L. built the experimental apparatus. A.L. and N.A. carried out experiments. A.L. and E.S. analyzed data and wrote the manuscript, with input from N.A.

## Declaration of interests

The authors have declared that no competing interests exist.

## Materials and Methods

### Experimental animals

A total of six adult hybrid mice were used in this study, one male and five females (**Table 1**). Hybrid mice were used since compared to the progenitors, hybrids exhibit reduced anxiety-like behavior, improved learning, and enhanced running behavior^21^. Four of the mice (m1-m4) were hybrid and double-transgenic, the F1 generation of an FVB/NJ female (JAX #001800, The Jackson Labs) and a male offspring of an Ai32 female (JAX #012569) and a CaMKII-Cre male (JAX #005359). The other two mice (m5 and m6) were offspring of an FVB/NJ female and a PV-Cre male (JAX #008069). In two subjects (m1 and m4), electrophysiological recordings and optical manipulations were carried out during some sessions. Results of electrophysiological recordings and optogenetic manipulations are not included in the present report. All behavioral results were observed at the subject level (**Table 1**; **Fig. S1**) and no differences were observed between implanted and un-implanted subjects. After separation from the parents, animals were housed in groups of same-litter siblings until participation in experiments. Animals were held on a reverse dark/light cycle (dark phase, from 8 AM until 8 PM). All animal handling procedures were in accordance with Directive 2010/63/EU of the European Parliament, complied with Israeli Animal Welfare Law (1994), and approved by the Tel Aviv University Institutional Animal Care and Use Committee (IACUC #01-16-051 and #01-21-061).

### Water deprivation

Mice were trained on a 2AFC task in which the rules governing discrimination behavior could change between different sessions. Every session was conducted on a different day. At the beginning of the training period, animals were housed one per cage and placed on a water-restriction schedule that guaranteed at least 40 ml/kg of water every day, corresponding to 1 ml for a 25 g mouse. Training was carried out five days a week, and animals received free water on the sixth day. Reward volume differed between mice and sessions, ranging 4-20 µl. The exact volume was determined by the experimenter before each session based on familiarity with the specific animal. In all sessions, the reward was larger by 20-50% during testing compared with training trials.

### Apparatus

The apparatus was a circular T-maze equipped with five motorized doors, five photosensors, two solenoid-driven reward ports, and a 100-LED visual stimulation matrix (**Fig. 1A**). All sensors and actuators were controlled by a microcontroller (Arduino Mega) via custom designed electronic circuitry. The home box (L x W x H: 20 x 30 x 10 cm) was located at the beginning of the central arm (75 x 8 x 3 cm) and connected to the end of the two lateral arms (100 x 6 x 3 cm). Each passageway between the home box and one of the arms was blocked by a transparent polycarbonate door. Two additional doors were located at the sides of the T-junction at the end of the central arm, blocking passage to the lateral arms. Every door was operated by a small motor (DC 6V 30RPM Gear Motor, Uxcell) and was equipped with two limit switches (D2F-01L2, Omron). There were five photosensors (PS; S51-PA-2-A00-NK, Datasensor). Water rewards were given by solenoids (003-0137-900, Parker), and each water port was connected to a different solenoid via flexible (Tygon) tubing. The visual stimulation matrix was constructed of 10 x 10 LEDs in alternating columns of blue (470 nm, Cree) and green (527 nm, Cree) diodes.

### Discrimination task

On a given session, a single rule including two associations was used (**Fig. 1D**). The allocation of rules to sessions was pseudo-random. Stimulus A was associated with leftward runs, and stimulus B was associated with rightward runs. Allocation of stimuli A and B to trials was pseudo-random. There were two types of sessions, “shaping” (pretraining) and “learning”. Shaping sessions included only “training” trials. Initially, each mouse was acquainted with the task in a series of shaping sessions (median [range]: 5 [4,10] sessions). Mice had to reach a criterion of 50 trials per session before commencing learning sessions. On each session, mice were free to perform the task until losing interest. Loss of interest was identified by behavior that included prolonged periods of rest and attempts to climb the walls of the home box. Learning sessions were divided into blocks (**Fig. 1B**), and every block included five training trials and ten “testing” trials.

A single “testing” trial proceeded as follows (**Fig. 1C**): (1) *Stimulus*: Once the animal entered the home box and passed the photosensors, the entrance doors (D4 and D5) closed and the exit door (D1) to the central arm opened (**Fig. 1A**). At the same time a stimulus A or B was given, remaining available to the animal until a lateral photosensor was passed during the response. (2) *Run*: Once the animal left the home box and passed the central arm photosensor, the two lateral doors (D2 and D3) at the T-junction opened, and the exit door (D1) closed. During the Run, the stimulus was continuously available. (3) *Response*: The animal chose a direction at the T-junction and went through one of the two open doors. Once the animal passed the lateral arm photosensor, the stimulus turned off, doors D2 and D3 closed preventing the animal from changing the choice, and doors D4 and D5 opened. (4) *Reward*: A liquid reward was available at the corresponding water port if the animal made a correct choice. The animal could not go back to the T-junction, but was free to consume the reward and return to the home box through the open home box door (D4 or D5).

During “training” trials, only the corresponding door (D2 or D3) at the T-junction opened. Because the animal could not make an incorrect response, a reward was given during every training trial.

### Quantification of rule difficulty and inter-rule similarity

To place arbitrary rules on a continuous scale, a symmetric intra-rule distance metric was introduced. Each stimulus was represented as a 100-element binary 2D array, with each element taking the value of 0 or 1 indicating whether the corresponding LED was off or on, ignoring the chromatic component. To derive the metric, every rule was characterized by three physical attributes that quantified the difference between the stimuli A and B. (1) The correlation distance (cd): One minus the maximal value of the 2D cross-correlation coefficient between A and B. (2) The Euclidean distance (ed): The scaled distance between the optimal match that yields the correlation distance. (3) The luminance distance (ld): The scaled difference in luminance between A and B. All distances (cd, ed, and ld) are non-negative scalars limited to the [0,1] range. The distance metric was then the magnitude of the 3D vector, defined as

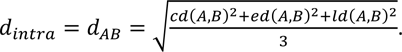

The metric ranges [0,1], taking the value of 0 when the two stimuli are identical (an impossibly-difficult rule) and 1 when stimuli are maximally distinct (a very easy rule).

The same physical attributes that characterize each rule were used to measure the difference between distinct rules (e.g., **Fig. 4D**). The inter-rule distance is a symmetric measure of the difference between the physical properties of the same-laterality stimuli of the two rules. Thus, for a pair of rules {A; B} and {A’; B’}, the inter-rule distance is the average of the intra-rule distances for two “mixed” rules: {A; A’} and {B; B’}. The resulting metric is

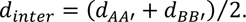

The metric takes the value of 0 when the two rules are identical and 1 when the rules are maximally distinct.

### Comparison of rank correlations

To determine if there is a difference between two rank correlation coefficients (cc_1_ and cc_2_), we bootstrapped (resampled with replacement) the two datasets (n=1000 iterations) and calculated cc_1_’ and cc_2_’ for each iteration. We then calculated the difference between the pairs of cc_1_’ and cc_2_’ and quantified the probability (two-tailed) that the difference differs from zero.

### Statistical analyses

In all statistical tests used in this study, a significance threshold of α=0.05 was used. All descriptive statistics (n, median, IQR, range, mean, SD, and SEM) can be found in the text, figures, and figure legends. Nonparametric statistical tests were used throughout. Differences between medians of two groups were tested with Mann-Whitney’s U-test (two-tailed). Differences between medians of three or more groups were tested with Kruskal-Wallis nonparametric analysis of variance and corrected for multiple comparisons using Tukey’s procedure. Wilcoxon’s signed-rank test was employed to determine whether a group median is distinct from zero (two-tailed). To estimate whether a given fraction was smaller or larger than expected by chance, an exact Binomial test was used. A permutation test was used to estimate significance of rank correlation coefficients (cc-s and pcc-s). For all figures, ns: p>0.05; *: p<0.05; **: p<0.01; ***: p<0.001.

## Supplementary figures

**Figure S1.**
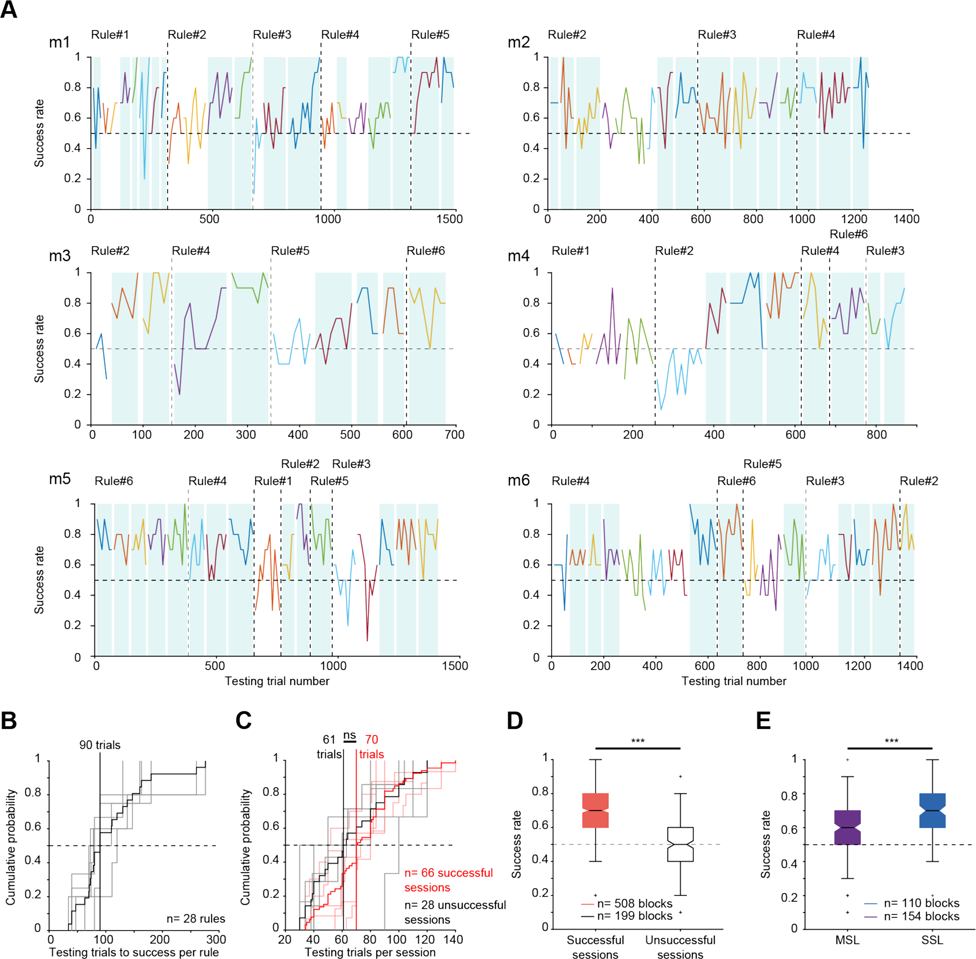
All mice achieve single session learning of at least one rule. (**A**) Every panel shows the success rate as a function of testing trial number for one subject. Sessions are represented by distinct colors, and every point indicates single-block success rate. A blue background highlights successful sessions (p<0.05, Binomial test comparing to chance level, 0.5). (**B**) Mice learn a new rule within a median of 90 testing trials. Data are from six mice tested on 28 rules. Grey lines, individual mice. (**C**) The number of testing trials performed during successful and unsuccessful sessions are not consistently different. Here and in **DE**, ns/***: p>0.05/p<0.001, U-test. (**D**) Success rates are higher during successful compared with unsuccessful sessions. (**E**) Success rates are higher during SSL compared with MSL rules.

**Figure S2.**
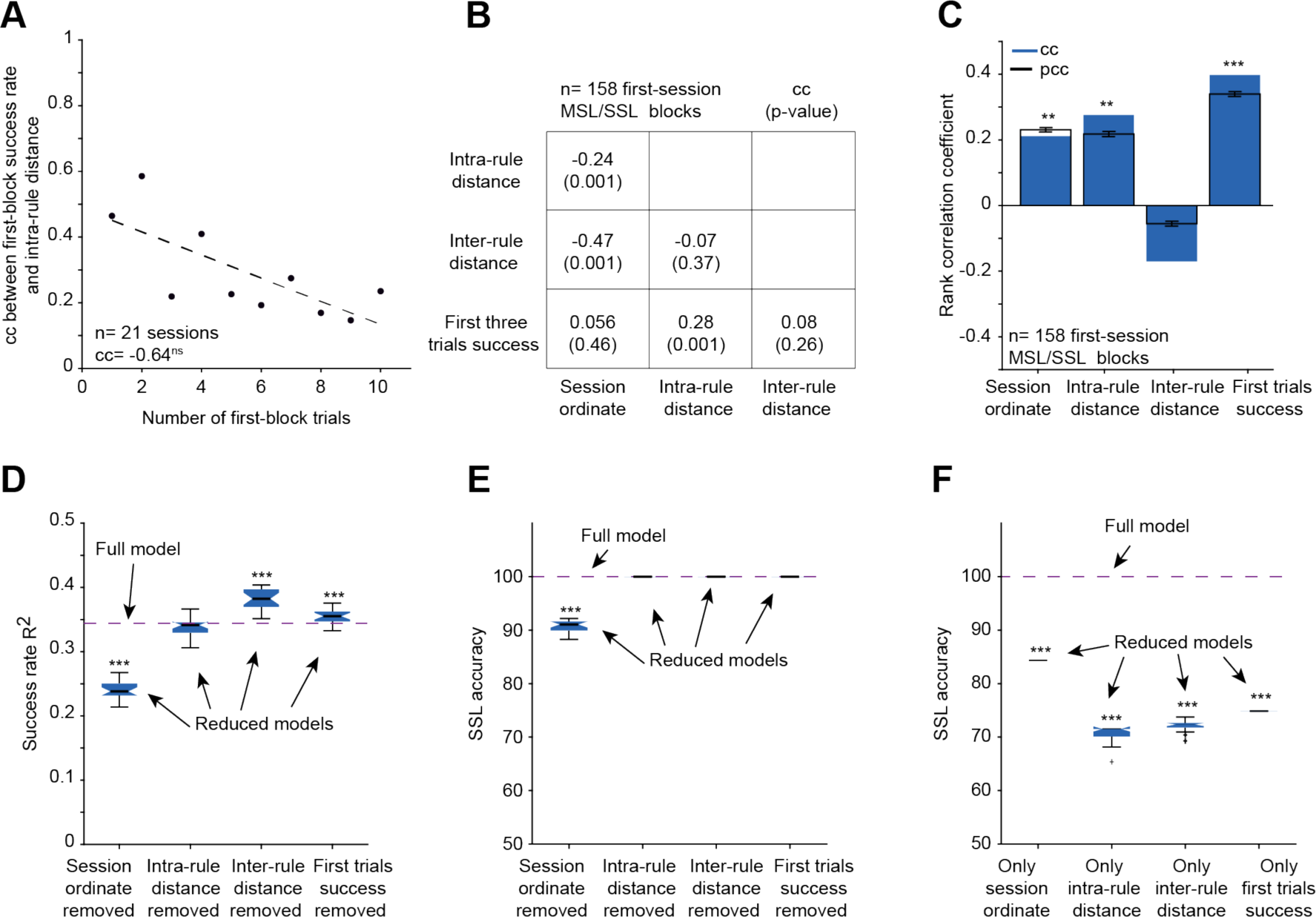
Success in the first three testing trials is correlated with rule difficulty. (**A**) Success rates during the first 1, 2, …, or 10 trials of SSL and MSL sessions is correlated with intra-rule distance. cc, rank correlation coefficient. Here and in **C**, ns/*/**/***: p>0.05/p<0.05/p<0.01/p<0.001, permutation test. (**B**) Correlation matrix for the three features used in **Fig. 4B-G** (session ordinate, intra-rule distance, inter-rule distance) and a fourth feature, the success rate during the first three trials. p-value, permutation test. (**C**) cc-s and pcc-s between success rate and the four features described in **B**. Error bars, SD. (**D**) Variance in block success rate (R^2^) explained by cross-validated support vector regression models. ***: p<0.001, Bonferroni-corrected U-test between the R^2^ of the full model and R^2^ of every reduced model, obtained by removing one feature. (**E**) Accuracy in predicting SSL using cross-validated support vector classification using the four-feature (full) model; and reduced three-feature models. Here and in **F**, ***: p<0.001, Bonferroni-corrected U-test between the accuracy of the full and reduced models. (**F**) Accuracy in predicting SSL using single-feature models.

## Notes

### Competing Interest Statement

The authors have declared no competing interest.

### Summary of Updates

All figures revised, table added, textual modifications throughout.

## References

1. Thorndike, E.L. (1927). The law of effect. Am. J. Psychol. 39, 212–222.

2. Carey, S., and Bartlett, E. (1978). Acquiring a single new word. Papers and Reports on Child Language Development. 15, 17–29.

3. Samborska, V., Butler, J.L., Walton, M.E., Behrens, T.E.J., and Akam, T. (2022). Complementary task representations in hippocampus and prefrontal cortex for generalizing the structure of problems. Nat. Neurosci. 25, 1314–1326.

4. Murphy, R.A., Mondragón, E., and Murphy, V.A. (2008). Rule learning by rats. Science 319, 1849– 1851.

5. Kurt, S., and Ehret, G. (2010). Auditory discrimination learning and knowledge transfer in mice depends on task difficulty. Proc. Natl. Acad. Sci. USA. 107, 8481–8485.

6. Broschard, M.B., Kim, J., Love, B.C., and Freeman, J.H. (2021). Category learning in rodents using touchscreen-based tasks. Genes. Brain. Behav. 20, e12665.

7. Reinert, S., Hübener, M., Bonhoeffer, T., and Goltstein, P.M. (2021). Mouse prefrontal cortex represents learned rules for categorization. Nature 593, 411–417.

8. Higgins, G.A., Grzelak, M.E., Pond, A.J., Cohen-Williams, M.E., Hodgson, R.A., and Varty, G.B. (2007). The effect of caffeine to increase reaction time in the rat during a test of attention is mediated through antagonism of adenosine A2A receptors. Behav. Brain Res. 185, 32–42.

9. Chun, M.M., and Jiang, Y. (1998). Contextual cueing: Implicit learning and memory of visual context guides spatial attention. Cogn. Psychol. 36, 28–71

10. Harlow, H.F. (1949). The formation of learning sets. Psychol. Rev. 56, 51–65.

11. Tse, D., Langston, R.F., Kakeyama, M., Bethus, I., Spooner, P.A., Wood, E.R., Witter, M P., and Morris, R.G. (2007). Schemas and memory consolidation. Science 316, 76–82.

12. Alonso, A., van der Meij, J., Tse, D., Genzel, L. (2020). Naïve to expert: Considering the role of previous knowledge in memory. Brain Neurosci. Adv. 4, 2398212820948686.

13. Bruce, H.M. (1959). An exteroceptive block to pregnancy in the mouse. Nature 184, 105.

14. Fanselow, M.S., and Bolles, R.C. (1979). Naloxone and shock-elicited freezing in the rat. J. Comp. Physiol. Psychol. 93, 736–744.

15. Cummings, K.A., and Clem, R.L. (2020). Prefrontal somatostatin interneurons encode fear memory. Nat. Neurosci. 23, 61–74.

16. Dudchenko, P.A. (2004). An overview of the tasks used to test working memory in rodents. Neurosci. Biobehav. Rev. 28, 699–709.

17. Rosenberg, M., Zhang, T., Perona, P., and Meister, M. (2021). Mice in a labyrinth show rapid learning, sudden insight, and efficient exploration. eLife 10, e66175.

18. Genzel, L., Schut, E., Schröder, T., Eichler, R., Khamassi, M., Gomez, A., Navarro Lobato, I., Battaglia, F. (2019). The object space task shows cumulative memory expression in both mice and rats. PLoS Biol. 17:e3000322.

19. Morris, R.G.M. (1981). Spatial localization does not require the presence of local cues. Learn. Motiv. 12, 239–260.

20. Morris, R. (1984). Developments of a water-maze procedure for studying spatial learning in the rat. J. Neurosci. Methods 11, 47–60.

21. Sloin, H.E., Bikovski, L., Levi, A., Amber-Vitos, O., Katz, T., Spivak, L., Someck, S., Gattegno, R., Sivroni, S., Sjulson, L., et al. (2022). Hybrid offspring of C57BL/6J mice exhibit improved properties for neurobehavioral research. eNeuro 9, ENEURO.0221-22.2022.

22. Burwell, R.D., Saddoris, M.P., Bucci, D.J., and Wiig, K.A. (2004). Corticohippocampal contributions to spatial and contextual learning. J. Neurosci. 24, 3826–3836.

23. O’Keefe, J., and Nadel, L. (1978). The hippocampus as a cognitive map. (Oxford University Press).

24. Carandini, M., and Churchland, A.K. (2013). Probing perceptual decisions in rodents. Nat. Neurosci. 16, 824–831.

25. Burgess, C.P., Lak, A., Steinmetz, N.A., Zatka-Haas, P., Bai Reddy, C., Jacobs, E.A.K., Linden, J.F., Paton, J.J., Ranson, A., Schröder, S., Soares, S., Wells, M.J., Wool, L.E., Harris, K.D., and Carandini, M. (2017). High-yield methods for accurate two-alternative visual psychophysics in head-fixed mice. Cell Rep. 20, 2513–2524.

26. International Brain Laboratory, Aguillon-Rodriguez, V., Angelaki, D., Bayer, H., Bonacchi, N., Carandini, M., Cazettes, F., Chapuis, G., Churchland, A.K., Dan, Y., Dewitt, E., Faulkner, M., Forrest, H., Haetzel, L., Häusser, M., Hofer, S.B., Hu, F., Khanal, A., Krasniak, C., Laranjeira, I., Mainen, Z.F., Meijer, G., Miska, N.J., Mrsic-Flogel, T.D., Murakami, M., Noel, J.P., Pan-Vazquez, A., Rossant, C., Sanders, J., Socha, K., Terry, R., Urai, A.E., Vergara, H., Wells, M., Wilson, C.J., Witten, I.B., Wool, L.E., and Zador, A.M. (2021). Standardized and reproducible measurement of decision-making in mice. eLife 10, e63711.

27. Marks, W.D., Osanai, H., Yamamoto, J., Ogawa, S.K., and Kitamura, T. (2019). Novel nose poke-based temporal discrimination tasks with concurrent in vivo calcium imaging in freely moving mice. Mol. Brain. 12, 90.

28. Yu, Y., Hira, R., Stirman, J.N., Yu, W., Smith, I.T., and Smith, S.L. (2018). Mice use robust and common strategies to discriminate natural scenes. Sci. Rep. 81, 1379.

29. Olton, D., Becker, J., and Handelmann, G. (1979). Hippocampus, space, and memory. Behav. Brain Sci. 2, 313–322.

30. Bernaud, V.E., Hiroi, R., Poisson, M.L., Castaneda, A.J., Kirshner, Z.Z., Gibbs, R.B., and Bimonte-Nelson, H.A. (2021). Age impacts the burden that reference memory imparts on an increasing working memory load and modifies relationships with cholinergic activity. Front. Behav. Neurosci. 15, 610078.

31. Bimonte-Nelson, H.A., Daniel, J.M., and Koebele, S. V. (2015). The mazes. In Theories, Practice, and Protocols for Testing Rodent Cognition, H.A. Bimonte-Nelson, ed. (Humana Press), pp. 37–72.

32. Schwegler, H., Crusio, W.E., and Brust, I. (1990). Hippocampal mossy fibers and radial-maze learning in the mouse: A correlation with spatial working memory but not with non-spatial reference memory. Neuroscience 34, 293–298.

33. Broadbent, N.J., Squire, L.R., and Clark, R.E. (2007). Rats depend on habit memory for discrimination learning and retention. Learn. Mem. 14, 145–151.

34. Seger, C.A., and Peterson, E.J. (2013). Categorization=decision making+generalization. Neurosci. Biobehav. Rev. 37, 1187–1200.

35. Shadmehr, R., and Mussa-Ivaldi, F.A. (1994). Adaptive task of dynamics during learning of a motor task. J. Neurosci. 14, 3208–3224.

36. Maes, E., De Filippo, G., Inkster, A.B., Lea, S.E.G., De Houwer, J., D’Hooge, R., Beckers, T., and Wills, A.J. (2015). Feature-versus rule-based generalization in rats, pigeons and humans. Anim. Cogn. 18, 1267.

37. Brooks, D.I., Ng, K.H., Buss, E.W., Marshall, A.T., Freeman, J.H., and Wasserman, E.A. (2013). Categorization of photographic images by rats using shape-based image dimensions. J. Exp. Psychol. Anim. Behav. Process. 39, 85–92.

38. Broschard, M.B., Kim, J., Love, B.C., Wasserman, E.A., and Freeman, J.H. (2019). Selective attention in rat visual category learning. Learn. Mem. 26, 84–92.

39. Levi, A., Spivak, L., Sloin, H.E., Someck, S., and Stark, E. (2022). Error correction and improved precision of spike timing in converging cortical networks. Cell Rep. 40, 111383.

40. Senzai, Y., and Scanziani, M. (2022). A cognitive process occurring during sleep is revealed by rapid eye movements. Science 377, 999–1004.

41. Sloin, H.E., Levi, A., Someck, S., Spivak, L., and Stark, E. (2022). High fidelity theta phase rolling of CA1 neurons. J Neurosci. 42, 3184–3196.

42. Voitov, I., and Mrsic-Flogel, T.D. (2022). Cortical feedback loops bind distributed representations of working memory. Nature 608, 381–389.

43. Rogers, S., Rozman, P.A., Valero, M., Doyle, W.K., and Buzsáki, G. (2021). Mechanisms and plasticity of chemogenically induced interneuronal suppression of principal cells. Proc. Natl. Acad. Sci. USA. 118, e2014157118.

44. Tarnavsky Eitan, A., Someck, S., Zajac, M., Socher, E., and Stark, E. (2021). Outan: An on-head system for driving µLED arrays implanted in freely moving mice. IEEE Trans. Biomed. Circuits Syst. 15, 303– 313.

45. Spivak L., Someck S., Levi A., Sivroni S., and Stark E. (2024) Wired together, change together: Spike timing modifies transmission in converging assemblies. Sci. Adv. in press.

46. Sadtler, P.T., Quick, K.M., Golub, M.D., Chase, S.M., Ryu, S.I., Tyler-Kabara, E.C., Yu, B.M., and Batista, A.P. (2014). Neural constraints on learning. Nature 512, 423–426.

47. Pakan, J.M.P., Francioni, V., and Rochefort, N.L. (2018). Action and learning shape the activity of neuronal circuits in the visual cortex. Curr. Opin. Neurobiol. 52, 88–97.

48. Poort, J., Khan, A.G., Pachitariu, M., Nemri, A., Orsolic, I., Krupic, J., Bauza, M., Sahani, M., Keller, G.B., Mrsic-Flogel, T.D., and Hofer, S.B. (2015). Learning enhances sensory and multiple non-sensory representations in primary visual cortex. Neuron 86, 1478–1490.

49. Jurjut, O., Georgieva, X.P., Busse, L., and Katzner, S. (2017). Learning enhances sensory processing in mouse V1 before Improving behavior. J. Neurosci. 27, 6460–6474.

50. Lak, A., Okun, M., Moss, M.M., Gurnani, H., Farrell, K., Wells, M.J., Reddy, C.B., Kepecs, A., Harris, K.D., and Carandini, M. (2020). Dopaminergic and prefrontal basis of learning from sensory confidence and reward value. Neuron 105, 700–711.

51. Le Merre, P., Esmaeili, V., Charrière, E., Galan, K., Salin, P.A., Petersen, C.C.H., and Crochet, S. (2018). Reward-based learning drives rapid sensory signals in medial prefrontal cortex and dorsal hippocampus necessary for goal-directed behavior. Neuron 97, 83–91.

52. Chevée, M., Finkel, E.A., Kim, S.J., O’Connor, D.H., and Brown, S.P. (2022). Neural activity in the mouse claustrum in a cross-modal sensory selection task. Neuron 110, 486–501.

53. Yang, G.R., Joglekar, M.R., Song, H.F., Newsome, W.T., and Wang, X.J. (2019). Task representations in neural networks trained to perform many cognitive tasks. Nat. Neurosci. 22, 297–306.

54. Molano-Mazón, M., Shao, Y., Duque, D., Yang, G.R., Ostojic, S., and de la Rocha, J. (2023). Recurrent networks endowed with structural priors explain suboptimal animal behavior. Curr. Biol. 33, 622–638.e7.

